# Altered transcription factor binding events predict personalized gene expression and confer insight into functional cis-regulatory variants

**DOI:** 10.1101/228155

**Authors:** Wenqiang Shi, Oriol Fornes, Wyeth W. Wasserman

**Affiliations:** Centre for Molecular Medicine and Therapeutics, Department of Medical Genetics, BC Children’s Hospital Research Institute, University of British Columbia, 950 28th Ave W, Vancouver, BC V5Z 4H4, Canada; Bioinformatics Graduate Program, University of British Columbia, 2329 West Mall, Vancouver, BC V6T 1Z4, Canada

## Abstract

Deciphering the functional roles of *cis*-regulatory variants is a critical challenge in genome analysis and interpretation. We hypothesize that altered transcription factor (TF) binding events are a central mechanism by which *cis*-regulatory variants impact gene expression. We present TF2Exp, the first gene-based framework (to our knowledge) to predict the impact of altered TF binding on personalized gene expression based on *cis*-regulatory variants. Using data from lymphoblastoid cell lines, TF2Exp models achieved suitable performance for 3,060 genes. Alterations within DNase I hypersensitive, CTCF-bound, and tissue-specific TF-bound regions were the greatest contributors to the models. Our *cis*-regulatory variant-based TF2Exp models performed as well as the state-of-the-art SNP-based models, both in cross-validation and external validation. In addition, unlike SNP-based models, our TF2Exp models have the unique advantages to evaluate impact of uncommon variants and distinguish the functional roles of variants in linkage disequilibrium, showing broader utility for future human genetic studies.

## Introduction

Understanding the functional role of genetic variants in human disease is a fundamental challenge in medical genetics. Whole genome sequencing now enables clinicians to systematically seek variants that contribute to disease phenotypes, but current clinical approaches focus primarily on the ~2% of the genome coding for proteins. Predicting the functional impact of non-coding variants remains a challenge, which limits interpretive capacity. As up to 88% of disease-related variants in genome-wide association studies (GWAS) are located within non-coding regions ^1^, there is a recognized need for methods that provide mechanistic insights into *cis*-regulatory variants.

Substantial progress has been made on detecting statistical relationships between common polymorphisms and expression levels. These expression quantitative trait loci (eQTL) studies can highlight regions harboring regulatory roles. Reported eQTLs are enriched for regulatory regions^2, 3^. Partially based on the success of eQTL analysis, regression-based models using SNPs proximal to genes as features have been developed, which show capacity to predict gene expression levels^4, 5^.

Such correlative approaches are useful, yet for multiple reasons they lack the resolution to direct researchers to specific causal alterations. First, causal variants are hard to infer in association studies due to linkage disequilibrium (LD) between SNPs ^6^. Second, uncommon variants (minor allele frequency, MAF < 0.05) are excluded from most association studies, but rare variants (MAF < 0.01) are often causal for genetic disorders^7, 8^. Third, most approaches defer the annotation of variant function until after the model is constructed, whereas an early focus on variants likely to impact gene regulation would provide more functional insight.

Both GWAS and eQTL studies have convincingly highlighted the importance of *cis*-regulatory regions^2, 3^. Advances in genomics and bioinformatics have greatly expanded the identification of functional elements within such regions, with an emphasis on DNA binding transcription factors (TFs). TFs recognize and bind to short DNA segments, named TF binding sites (TFBSs), in a sequence-specific manner ^9^. Machine learning approaches coupled to extensive TF ChIP-seq data have enabled better predictions of TFBSs^10, 11^. Recently, the compilation of altered TF binding events has increased, and models have emerged to predict such events^12, 13^. However, the relationship between altered TF binding events and gene expression levels remains unclear, hindering our understanding of *cis*-regulatory variants^14, 15^.

To gain more direct insight into the functional roles of non-coding variants, a key challenge is to determine the relationships between alterations of TF binding events and observed expression levels of a target gene. To address this challenge, we developed TF2Exp models to predict gene expression levels based on TF binding alterations inferred from *cis*-regulatory variants. We explored the utility of TF2Exp in answering four important questions: 1) are alterations of TF binding events predictive of gene expression changes?; 2) what are the characteristics of the functional altered TF binding events?; 3) do TF2Exp models perform as well as the state-of-the-art SNP-based models?; and 4) are TF2Exp models able to evaluate the impact of SNPs in LD and uncommon variants? Our results show that TF2Exp models successfully predicted the alteration of gene expression for over three thousand genes, with an average performance comparable to that of models based solely on SNPs, supporting the hypothesis that TF binding alteration is a central mechanism by which *cis*-regulatory variants impact gene expression.

## Results

### TF2Exp: regression models to predict the impact of altered TF binding on gene expression

We developed TF2Exp, a gene-based computational framework to assess the impact of altered TF binding events on gene expression (Figure 1). As detailed in Materials and methods, variant calling data (single nucleotide variants and small indels) and gene expression data for 358 lymphoblastoid cell lines (LCLs) were obtained from the 1000 Genomes ^16^ and GEUVADIS projects ^3^. Moreover, TF-bound regions for 78 distinct TFs and DNase I hypersensitivity sites (DHSs) were obtained from the ENCODE project for GM12878 LCL ^2^. TF binding events (inclusive of DHS) were associated to a gene if they overlapped either the promoter or distal regulatory region of the gene (see Materials and methods). The impact of each single variant within a TF binding event was scored using DeepSEA ^10^, and multiple variants within the same TF binding event were summed to generate an overall alteration score of that TF binding event in each individual. On average, each gene had 420.0 altered TF binding events within 36.6 regulatory regions across the 358 individuals. Based on computed alteration scores of TF binding events in each individual, regression models were trained by LASSO^17^ to predict gene expression per individual and to identify key contributing TF binding events.

**Figure 1.**
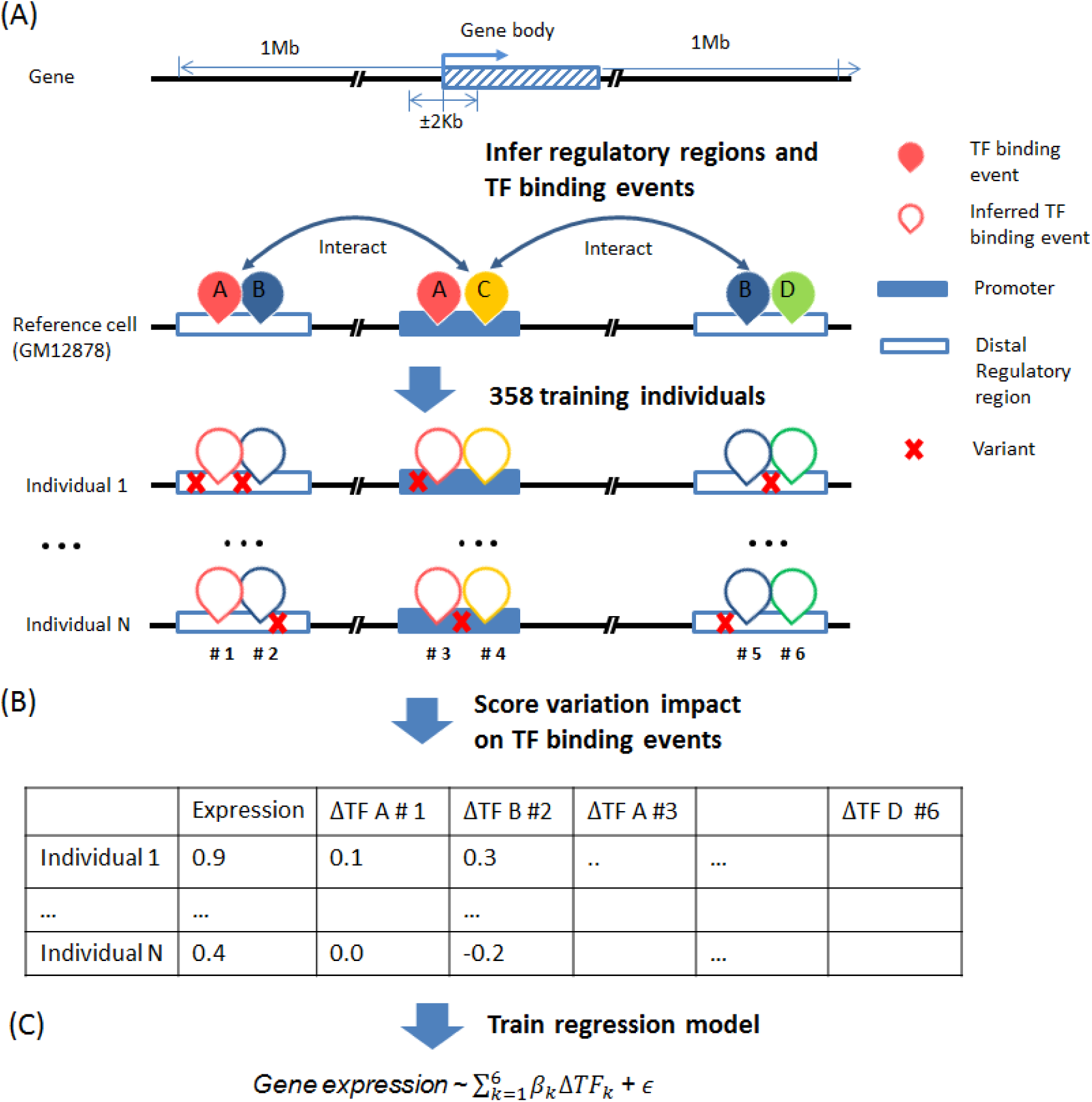
Overview of the TF2Exp framework. (A) Infer regulatory regions and TF binding events of each gene based on the reference cell line (GM12878). Distal regulatory regions are associated to a gene according to Hi-C data. TF binding events on the promoter or distal regulatory regions of a gene are assigned to that gene. (B) Score the alteration of TF binding events based on the overlapped variants for each individual. (C) Train regression models for each gene across the collected individuals.

### TF2Exp predicts the expression levels for a subset of genes

We successfully trained TF2Exp models for 15,914 genes. Average model performance (R^2^) by 10-fold cross-validation was 0.048, with most models having low predictive power (Figure 2). To focus on predictive models, we applied an R^2^ threshold of 0.05 as in ^4^, resulting in 19.2% of genes (hereinafter referred to as predictable genes). To assess the impact of random noise in the model training process, we set up control models in which gene expression was shuffled across individuals while preserving TF binding features. Control models achieved an average R^2^ of only 1.9×10^−4^ (Figure 2), supporting the non-random signal captured by TF2Exp models. As in the work of Manor *et al*. ^4^, we observed a significant correlation between model performance and the variance of expression levels for the predictable genes (Spearman correlation 0.25, p-value = 4.0×10^−43^; Supplementary file: Figure S2). We performed gene ontology enrichment analysis using GREAT ^18^. The top 10% predictable genes are enriched in pathways including graft-versus-host disease, allograft rejection and autoimmune thyroid disease, relevant to the roles of B cells (original cell type before transformed to LCL) in the immune system.

We next sought to determine if additional information could substantially improve model performance. We assessed whether prior knowledge, such as Hi-C proximity scores and known TF-TF physical interactions, could improve TF2Exp models. We introduced the proximity scores of Hi-C interactions to guide model fitting, so that TF-binding events on highly-interacting regions would be less regularized by LASSO (Materials and methods). We observed that adding Hi-C proximity scores resulted in a slight R^2^ improvement of 9.4×10^−4^ (Wilcoxon signed-rank test, p-value = 8.1×10^−45^), suggesting that the original TF2Exp models had captured most of the signal from the Hi-C data. We also tested models including interaction terms for known TF-TF physical interactions (Materials and methods). Adding TF-TF interactions significantly reduced model performance by 7.7×10^−4^ (Wilcoxon signed-rank test, p-value = 2.2×10^−81^, Figure 2), suggesting that TF-TF interaction terms did not add further information. Taken together, models incorporating prior knowledge achieved similar performance to the original models. Thus, we focused on the original (and simpler) TF2Exp models in the next stages of the analysis.

**Figure 2.**
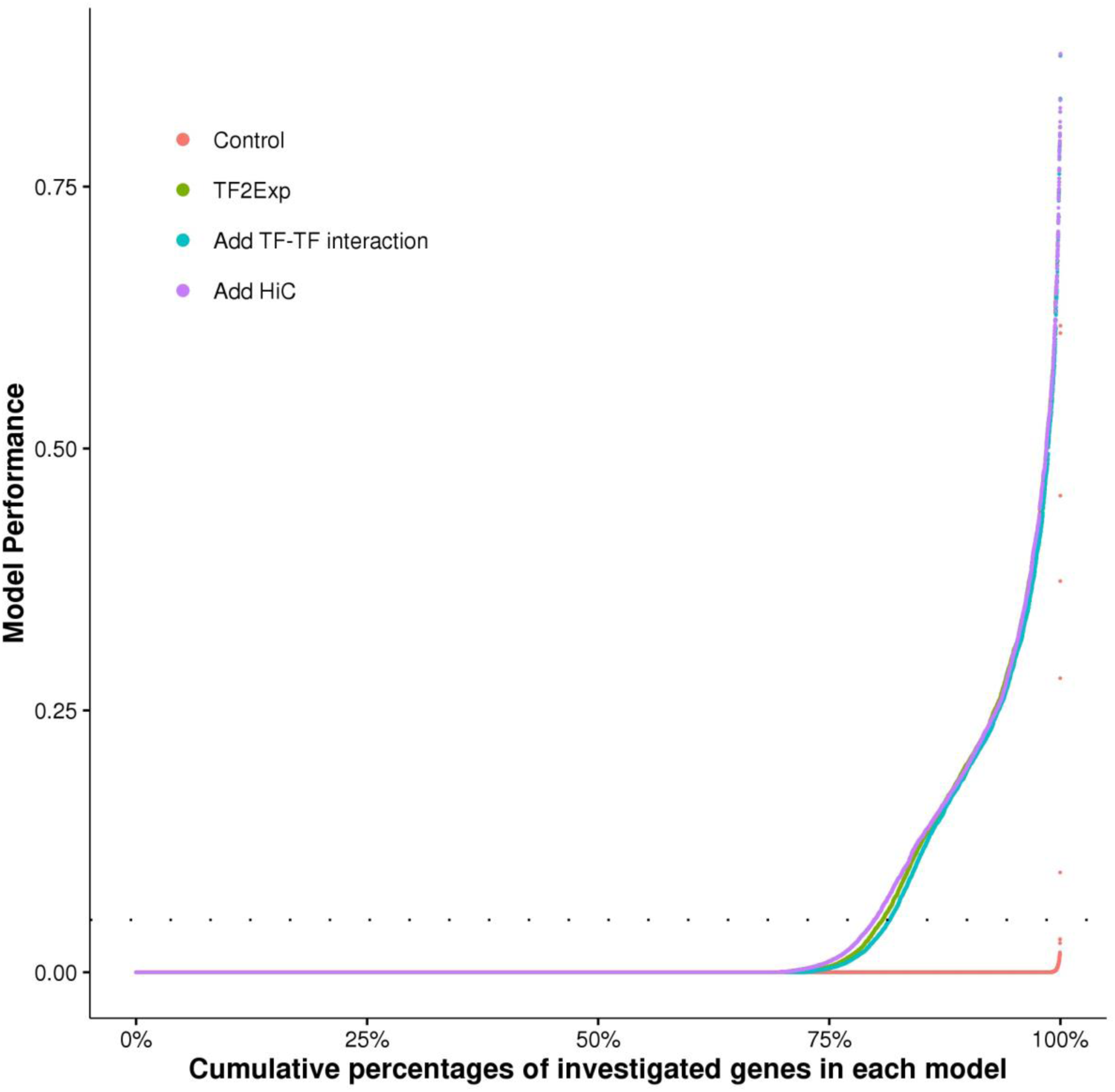
Performance comparison of alternative TF2Exp models. For each type of TF2Exp model, performances (R^2^) of investigated genes (y axis) are plotted in ascending order with respect to the cumulative percentage of genes (x axis). Dashed line indicates the defined performance threshold of 0.05 for predictable genes.

### Alterations of DHS, CTCF and tissue-specific TF binding are the most frequently selected features

We next sought to identify TFs for which binding events were more frequently selected in TF2Exp models. For the predictable genes, models selected an average of 3.7 key features (where a feature was the alteration score of a single TF binding event). Frequently selected TFs tended to have more binding events across the genome (Pearson correlation 0.97, p-value < 2.2×10^−16^). The top 5 selected TF features included DHS, RUNX3, CTCF, EBF1 and PU.1, accounting for 34.2% of the selected features (Figure 3). Particularly, 41.4% of the predictable genes had at least one DHS feature, highlighting the well-known relationship between chromatin accessibility and gene expression ^19^. CTCF has diverse roles in gene regulation across multiple tissues^20, 21^, and the remaining three TFs perform important roles in LCL tissue-specific regulation: RUNX3 in immunity and inflammation ^22^, EBF1 in B lymphocyte transcriptional network expression ^23^, and PU.1 in lymphoid development ^24^.

**Figure 3.**
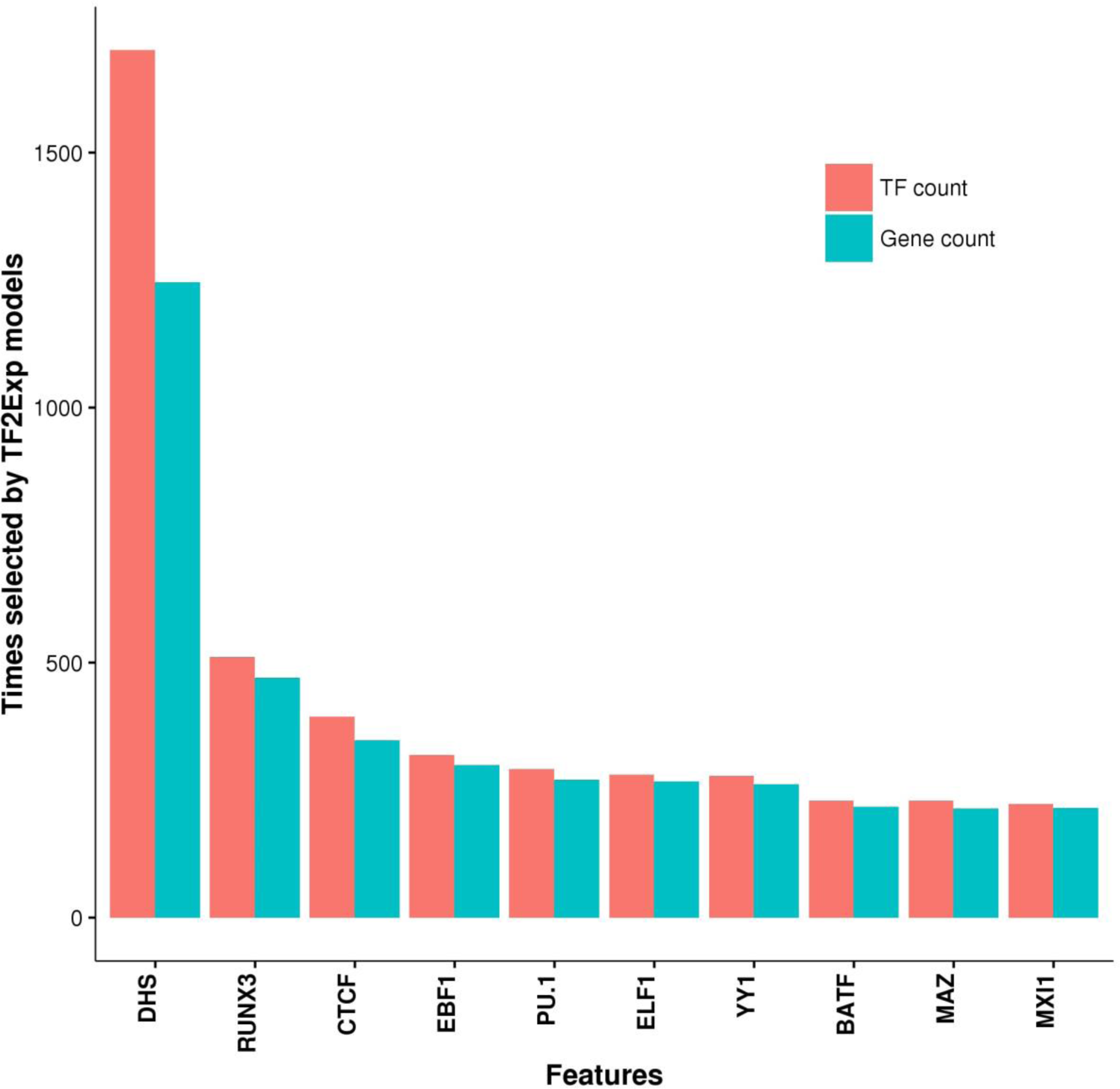
Top 10 TFs whose binding events are the most frequently selected features across predictable genes. Red bars indicate the total number of TF binding events selected by TF2Exp models. Blue bars indicate the total number of genes that selected binding events of the indicated TF as key features.

### Selected TF binding events correlate with gene expression in *vivo*

We next sought to assess whether *in vivo* TF binding of selected features correlated with gene expression. We obtained CTCF and PU.1 ChIP-seq LCL data for two independent sets of 45 originally training individuals (38 individuals overlapped between the two sets). TF binding signals were extracted from the reference GM12878 TF binding events (*i.e.* the ChIP-seq features used in the TF2Exp for model construction). In predictable genes, 83 CTCF and 72 PU.1 binding events were selected for testing based on their high variance of binding score change (see Materials and methods). Eight CTCF (9.7%) and seven PU.1 (9.6%) of the tested *in vivo* binding events significantly correlated with gene expression levels (Pearson correlation, FDR<0.05), and their correlation coefficients were consistent with the coefficients estimated based on the TF sequence alteration score and gene expression (p-value= 1.4×10^−4^, coefficient = 0.81). Due to limited testing sample size (n = 45), we did not have sufficient statistical power to detect weakly correlated TF-gene relationships (e.g. coefficient < 0.29, see Materials and methods), which accounted for most (89.7%) of the tested *in vivo* binding events. In summary, we observed that 9.7% of TF binding events selected by TF2Exp displayed detectable correlation (correlation coefficient > 0.29) between *in vivo* binding and gene expression.

### Effect sizes of TF binding events within promoters are greater than distal regulatory regions

We next examined the locations and effect sizes of selected features. The selected features in promoters were mostly within 10Kb of gene start positions, while selected features in distal regions were distributed within ~500Kb. We observed significant depletion of selected features in distal regulatory regions compared with promoter regions (Fisher’s exact test, odds ratio = 0.32, p-value < 2.2×10^−16^). Effect sizes of TF binding events decreased rapidly in relation to the distance from gene start positions (Figure 4A). Such a decreasing trend has been reported for effect sizes of eQTLs ^25^. The selected features in promoter regions also exhibited significantly larger absolute effect sizes (Wilcoxon rank-sum test, p-value = 2.5×10^−53^, Figure 4B) and more positive effects (Wilcoxon rank-sum test, p-value = 3.0×10^−4^) than features in distal regulatory regions. Nevertheless, the selected distal features of a gene were significantly enriched in the enhancer regions associated to that gene, as specified in the FANTOM5 project ^26^ (Fisher’s exact test, odds ratio = 1.3, p-value < 1.5×10^−9^, see Materials and methods), supporting a functional role of the selected distal TF binding events. Thus TF2Exp models are identifying *cis*-regulatory sequence variants that bring functional insights into the mechanisms underlying gene expression levels.

**Figure 4.**
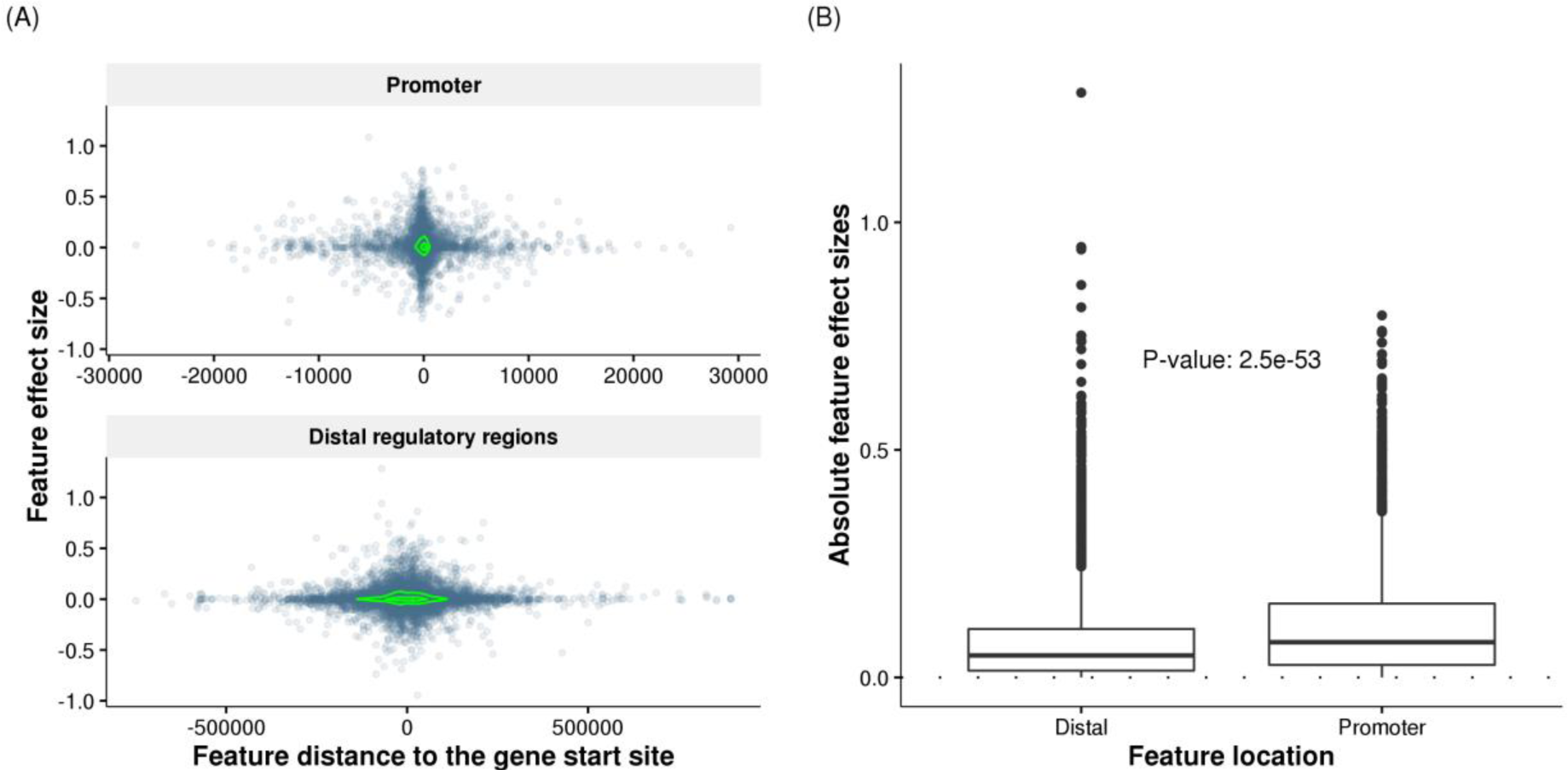
Feature effect sizes in promoter and distal regulatory regions. (A) Effect sizes of selected features decrease rapidly with their increasing distances to the gene start positions. Each dot represents one selected feature (TF binding event) of predictable genes, and the coordinates indicate the feature distance to gene start site (x axis) and the feature effect size (y axis) obtained in TF2Exp models. The green contours indicate estimated dot density.

Feature effect sizes are plotted separately for promoter regions (top panel) and distal regulatory regions (bottom panel). (B) Compare the absolute feature effect sizes of selected TF-binding events at promoters and distal regulatory regions across the all the predictable genes. The labeled p-value indicates the significance for the difference of two groups (Wilcoxon rank-sum test).

### Uncommon variants improve model performance for a small portion of genes

As TF2Exp models can distinguish the impact of variants in TF-binding events, we investigated the contribution of uncommon (MAF <= 0.05) variants to model performance. TF2Exp models trained only on uncommon variants achieved lower average performance (R^2^ = 0.011) compared with models based on all variants (R^2^ = 0.048). However, when combining both uncommon and common variants, a small portion (11.5%) of models improved compared with using common variants alone. The improvement can be negative if performances of uncommon variants models were near zero (Supplementary file: Figure S3), suggesting that majority of the uncommon variants are not informative for TF2Exp models.

### TF2Exp models can distinguish SNPs in LD compared with SNP-based expression models

We compared our TF2Exp models with state-of-the-art models, which predict alteration of gene expression levels using proximal SNPs^4, 5^ (see Materials and methods). First, for each gene, we trained both models (TF2Exp and SNP-based) on the same set of variants (SNPs within all TF binding events, SNPinTF) for each gene, and named two models as TF2Exp-SNPinTF and SNP-SNPinTF. The two models showed comparable performance across the shared predictable genes (Wilcoxon signed-rank test, p-value=0.19; Supplementary file: Figure S4). In addition, the default SNP-based models using all the proximal SNPs within 1Mb of the gene body showed essentially equal performance (mean R^2^ = 0.051) to TF-SNPinTF (mean R^2^ = 0.050, Wilcoxon signed-rank test, p-value=0.08), indicating that altered *cis*-regulatory variants can serve equally as well as SNPs as the basis for predictive expression models, while providing added benefit of mechanistic insight.

Compared with SNP-based models, TF2Exp models are able to infer the functional roles of SNPs in linkage disequilibrium (LD) based on the predicted impact of variants on TF-bound regions. Most of the selected SNPs (59.8%, n=9,386) in the SNP-SNPinTF models overlapped selected TF binding events (62.7%, n=12,663) in TF2Exp-SNPinTF for the same gene. 18.4% of the overlapped SNPs were in high LD (r^2^>0.9) with other SNPs in the same TF-bound regions, hindering the inference of the casual variants for SNP-based models. Based on TF2Exp models, we found that 36.8% of the linked SNPs showed at least a two-fold impact on the overlapped TF-bound region compared with the selected SNPs (Figure 5), suggesting a more dominant contribution of the linked SNPs. In addition, a subset of the selected SNPs (20.1%) overlapped more than one selected TF binding event, which highlights that individual SNPs can alter multiple mechanisms of gene regulation. Overall, TF2Exp models provide a quantitative way to evaluate the impact of SNPs in LD, suggesting a broader utility for genomic studies than SNP-based models.

**Figure 5.**
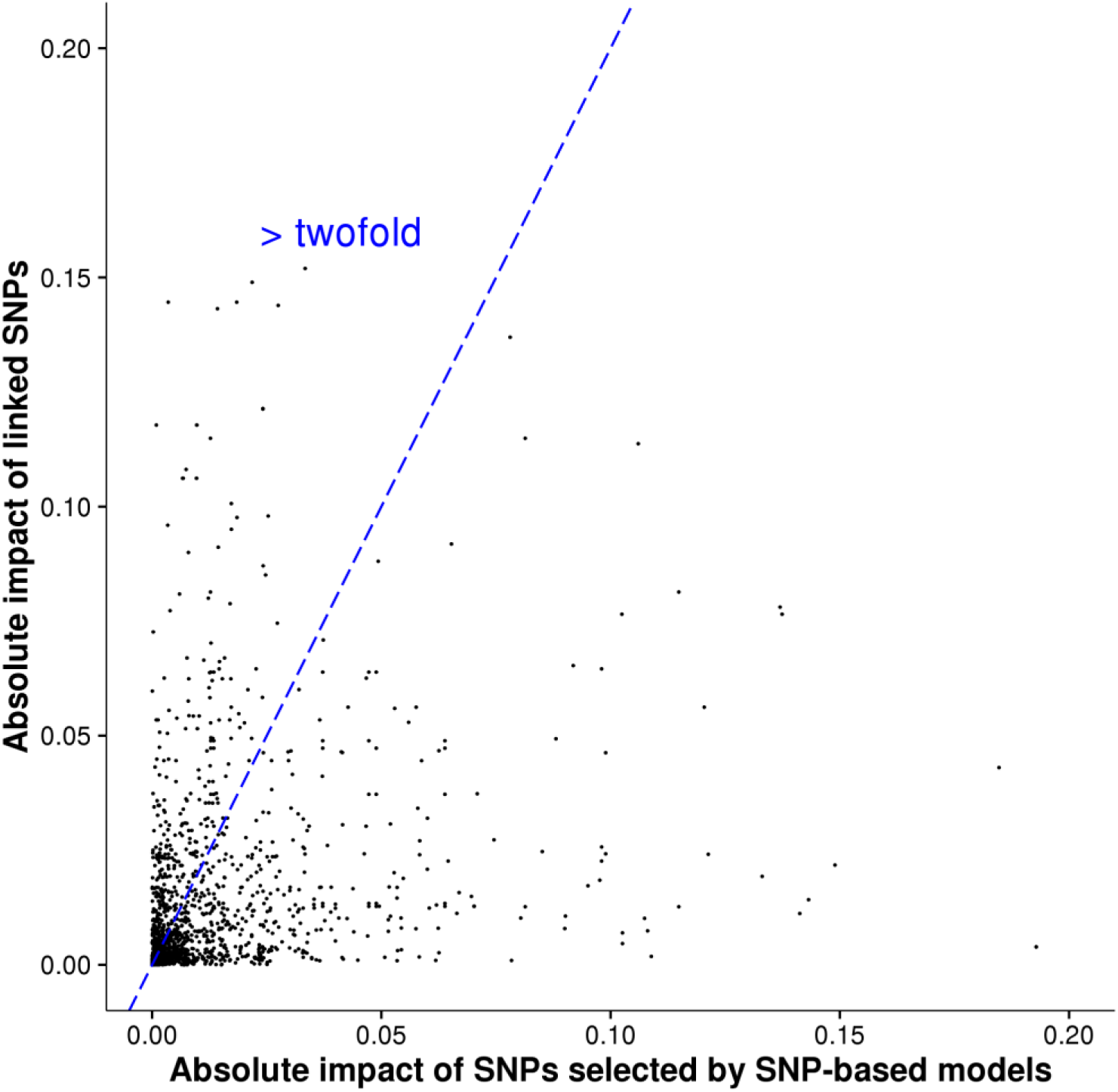
Distinguish the functional roles of SNPs in LD based on TF2Exp framework. Most of the SNPs selected by SNP-based models overlapped with TF binding events selected by TF2Exp models for the same gene. A subset of these selected SNPs were in high LD (r^2^>0.9) with other SNPs in the same TF-bound region. Each dot depicts the absolute impact of a selected SNP by a SNP-based model (x axis) versus the absolute impact of its linked SNP, according to the TF2Exp model. The dashed line indicates two-fold impact of the linked SNPs compared with the selected SNPs.

### TF2Exp models exhibit robust performance in external validation datasets

We finally sought to evaluate the models of predictable genes on external datasets. We obtained microarray expression data from LCLs of 256 individuals ^27^, including 80 Utah residents with Northern and Western European ancestry (CEU), 87 Chinese (CHB) and 89 Japanese (JPT) (Materials and methods). As 79 of the CEU individuals overlapped with the training individuals of TF2Exp models, we first evaluated the agreement between the microarray and RNA-seq data on these individuals. Relative expression levels across all genes within each individual were concordant between microarray and RNA-seq experiments (average Spearman correlation = 0.76), supporting an overall consistency between the two data sets. However, when we considered a single gene across the 79 individuals, the correlation between the two platforms was low (average Spearman correlation = 0.19). Therefore, we expected models trained on RNA-seq data to have an upper limit performance when applied to microarray data. Then, we used TF2Exp models to predict gene expression levels on the CHB and JPT individuals. TF2Exp models achieved an average correlation of 0.16 for CHB and 0.15 for JPT individuals. Similarly, SNP-based models achieved an average correlation of 0.16 for both populations.

An example of a high performing gene (FAM105A) in the external validation is illustrated in Figure 6A by comparing the predicted (TF2Exp) and observed (microarray) expression levels. FAM105A is associated with pancreatic islet function and type 2 diabetes^28, 29^. For this gene, TF2Exp identified 4 contributing TF binding events (Figure 6B), of which two of them had greater weights: DHS (chr22:45711760-45711910, effect size: -0.325) and MEF2A (chr22:45771822-45772122, effect size: 0.334). Alterations of these key events largely explained the changes of gene expression in the different individuals. For example, NA18640 had the lowest observed expression level in CHB individuals, as variant rs104664 of this individual was predicted by TF2Exp to increase the score of DHS; while rs5765304 in NA18573 increased MEF2A binding scores, resulting in the highest predicted expression.

**Figure 6.**
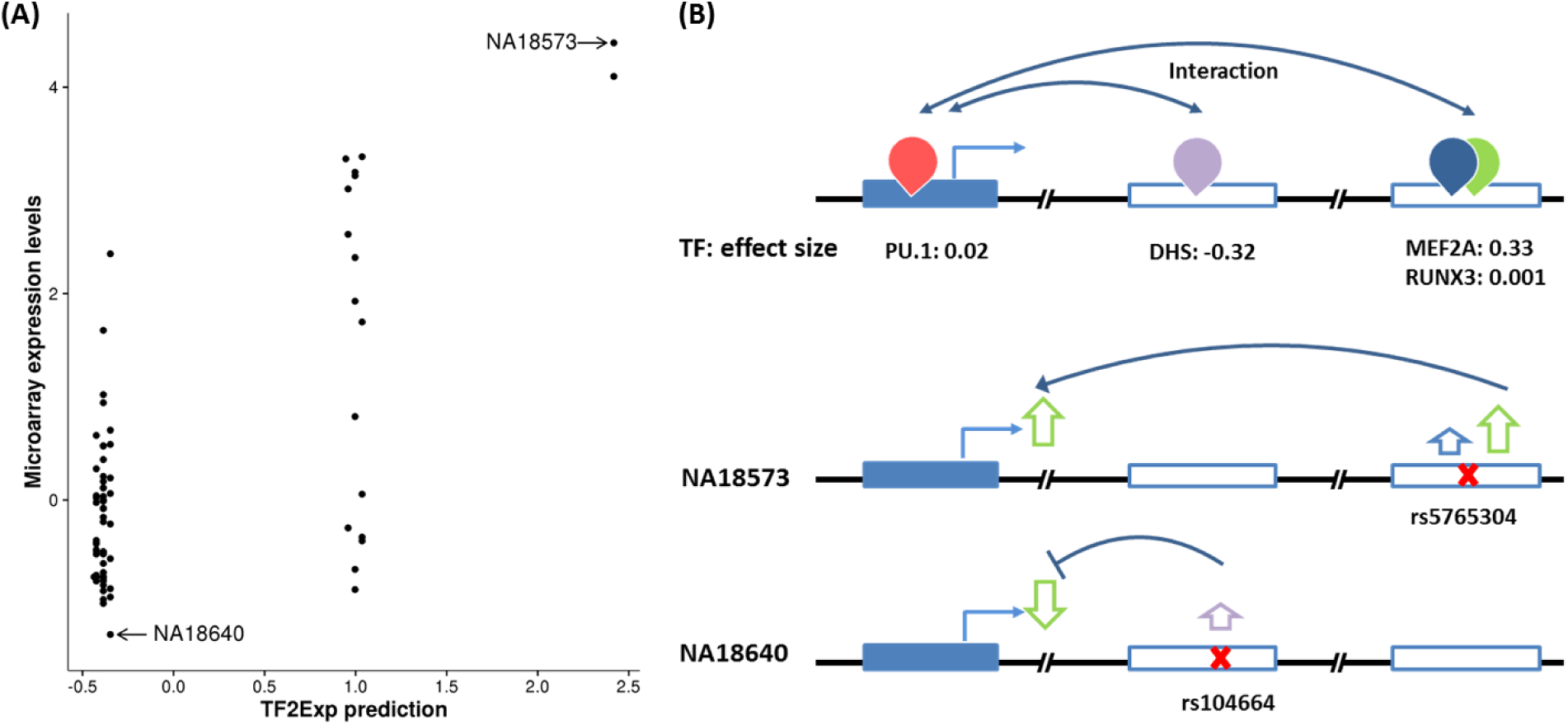
Performance and key features of TF2Exp for FAM105A gene in the external validation set. (A) Each point represents one tested individual and its coordinates indicate the predicted expression levels given by TF2Exp model (x axis) and the observed microarray expression (y axis). (B) Key features and inferred roles of variants of two individuals. The top panel illustrate key TF binding events learned from training data sets. The figure legend is the same as Figure 1. The middle and bottom panel show the variants within key TF binding events and their inferred roles on TF binding and gene expression for two individuals.

## Discussion

Deciphering the functional roles of regulatory variants is a critical challenge in the post-sequencing era. To address this challenge, we have introduced a novel framework, TF2Exp, which uses alteration of TF binding as an intermediate feature to elucidate the functional impact of regulatory variants and predict gene expression levels. TF2Exp models based on lymphoblastoid cell line data showed predictive capacity for over 3,000 genes, incorporating an average of 3.7 altered TF binding events per gene model. The most frequently selected TF binding events included both general properties (*e.g.* alterations within DNase I hypersensitive regions) and tissue-specific properties (*e.g.* alterations in TF bound regions for TFs relevant to the studied lymphoblastoid samples). TF2Exp models achieved equivalent performance to state-of-the-art SNP-based models, and provide mechanistic insights into *cis*-regulatory variants.

TF2Exp models have the potential to address two challenges left unresolved by SNP-based models and classical eQTL studies. For these approaches, it is difficult to: 1) infer variant function, as the studied SNPs can be in high linkage disequilibrium; and 2) evaluate the impact of rare variants (which are excluded from such analyses). By treating TF binding events as functional units within genes, TF2Exp models can evaluate the relative impact of any variant (SNV or small indel) within a TF-bound region. As in the example presented in Figure 6, for individual variants, the derived impact within the model is independent of the linkage disequilibrium or allele frequency. Moreover, even though the inclusion of uncommon variants only improved the performance for a small portion of genes (~11%), the resulting TF2Exp models offer a unique advantage for the inference of functional *cis*-regulatory variants, compared with previous SNP-based methods^4, 5^.

Similarly to SNP-based methods, the predictive performance of TF2Exp models is limited, showing utility for a subset of genes (19.2%), and even within these genes, model performance is modest (R^2^ = 0.21). The limited performance is likely attributable to multiple causes. First, the variance of gene expression attributed to common variants is quite low (*e.g.* 15.3% as estimated by Gamazon *et al.* ^5^), suggesting that models restricted to DNA sequence features alone only account for a portion of the observed variance in expression levels. Second, TF2Exp models were limited by the availability of ChIP-seq data (78 TFs in LCLs), while transcriptome studies have revealed that human cells express around 430 TFs on average ^30^. Third, TF2Exp models focused on TF binding events potentially involved in transcriptional regulation, but other regulatory mechanisms (*e.g.* post-transcriptional regulation) or genomic features (*e.g.* DNA methylation ^31^ or sequence conservation ^32^) might explain additional portion of the observed variance of gene expression. Fourth, TF2Exp models were likely constrained by the small number of available training samples, as including additional features (*e.g.* TF-TF interactions and uncommon variants) decreased model performance. We expect that the expansion of reference transcriptome datasets will provide more samples for exploring more complex relationships between genes and TF binding events, thereby improving model performance. Fifth, long-distance interactions within the nucleus ^33^ are unaccounted for in existing models, and incorporating more dimensions in the nucleus could further improve model performance.

In conclusion, identifying the impact of *cis*-regulatory variants on gene expression is a critical step towards understanding the genetic mechanisms contributing to diseases. TF2Exp models are able to predict the impact of altered TF binding on gene expression levels and provide mechanistic insights into the roles of selected TF-binding events and *cis*-regulatory variants. We anticipate that future enlarged omics data, in LCLs and other cell types, will greatly expand the application scope of TF2Exp models.

## Materials and methods

### Quantifying gene expression from RNA-seq data

LCL RNA-seq and variant calling data for 358 individuals from European populations were downloaded from the GEUVADIS project ^3^ and the 1000 Genomes Project ^34^ (Supplementary notes). Individuals covered 4 populations, including 89 Utah residents with Northern and Western European ancestry (CEU), 92 Finns (FIN), 86 British (GBR) and 91 Toscani (TSI). For each population, we built sex-specific transcriptomes in which SNP positions with MAF ≥ 0.05 were replaced by N (representing any of the four nucleotides A, C, G, T) using scripts from ^35^. RNA-seq data were processed using Sailfish (version 0.6.3) ^36^, and the expression level of each gene was quantified as transcripts per million reads. The resulting expression data were normalized via multiple steps, including standardization, variation stabilization, quantile normalization and batch effects removal (*i.e.* population and gender, and 22 hidden covariates) by PEER ^37^ (Supplementary file: Figure S1). Any gene that was either on the sex chromosomes or showed near-zero variance in expression levels was removed, leaving 16,354 genes for model training.

### Associating TF binding events to genes according to Hi-C data

We obtained Hi-C data to measure physical interactions between DNA regions (Hi-C fragments) from GM12878 cells (an LCL) ^35^. The average size of Hi-C fragments was 3.7Kb ^35^. Promoters were defined as the ±2Kb region centered at the start position of a gene (outermost transcript start position annotated by Ensembl ^38^ in genome assembly GRCh37). Each promoter was extended to include any overlapping Hi-C fragments. Distal regulatory regions were defined as Hi-C fragments within 1Mb of a gene body (as delimited by the outermost transcript start and end) interacting with the promoter of that gene (proximity score >0.4). Uniformly processed GM12878 DHSs and ChIP-seq peaks for 78 TFs were downloaded from the ENCODE project ^2^. As DHS is a general indicator of TF binding ^39^, DHSs are referred to as part of the set of ChIP-seq peaks within this manuscript for editorial convenience. A TF binding event was associated to a gene if it overlapped the promoter or a distal regulatory region of that gene. The resulting associations between genes and TF binding events derived from GM12878 cells were used as the reference for all studied individuals.

### Predicting sequence variation impact on TF binding events

Variant calling data of each individual was downloaded from the 1000 Genomes Project (release 20130502) ^34^. We only considered single nucleotide variants and small indels (<100bp). For each individual, the impact of a variant within a TF binding event was evaluated as the binding score difference between the altered and reference alleles, as determined by the corresponding DeepSEA (v0.93) TF binding model trained on GM12878 data ^10^. To allow for the analysis of multiple variants within a TF binding event, we modified DeepSEA to calculate the binding score of each allele using the 1,100bp region centered at the ChIP-seq peak max position (the original code centered the 1,100bp region at each variant). Score differences of multiple variants within the same TF binding event were summed to represent the overall alteration of that event. TF ChIP-seq peaks with multiple peak max positions and overlapped peaks from the same experiment were split at the center of each pair of neighboring peak max positions. At heterozygous positions, the binding score difference was divided by 2. Lastly, we calculated the linkage disequilibrium between variants across studies individuals using plink2 ^40^.

### Quantitative models of gene expression

#### LASSO regression on gene expression

We developed regression models to predict the expression level of a gene using altered TF binding events associated with that gene based on the following equation:

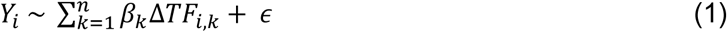

where *Y_i_* is the expression levels of gene *i* across the studied individuals, *n* is the number of TF binding events associated with gene *i*, Δ*TF_i,k_* is the alteration of TF binding event *k* across studied individuals and β_k_ is the effect size of TF binding event k. In equation (1), *Y_i_* is the response and Δ*TF_i,k_* is the input feature for the LASSO regression model, which was trained using the R ^41^ glmnet package ^17^ based on collected training data for 358 LCLs. Model performance was evaluated by 10-fold nested cross-validation, in which internal folds identified the optimal hyper-parameter lambda, and outer layers tested the model performance. Model performance was measured as the square of the correlation between predicted and observed expression levels (R^2^). The trained models would select a subset of TF binding events as key features of which effect sizes were not zero. When Hi-C proximity scores were used as the prior to select features, the prior (penalty.factor in the glmnet function) was set to “1 – proximity score”.

#### Defining TF-TF interactions

For TFs known to interact in the BioGrid database ^42^, we created interaction terms between pairs of TF binding events (one from each TF) if they satisfied one of the following conditions: 1) two binding events overlapped by at least 200bp; or 2) their regulatory regions were reported to interact in the Hi-C data.

#### SNP based models

For each gene, we trained regression models based on multiple SNPs to predict the expression level of that gene following the same procedure as in the work of Gamazon *et al.* ^5^. We only considered SNPs with MAF > 0.05 and within 1Mb of gene body regions. The regression formula for SNP-based models was as follows:

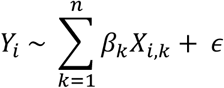

Where *Y_i_* is the expression levels of gene *i* across studied individuals, *n* is the number of SNPs, and *X_i,k_* is the number of minor alleles of *SNP_i,k_*.

### Analyzing selected features using FANTOM5 data

In the FANTOM5 project, an enhancer is associated with a gene based on the correlation of expression between the enhancer and the gene promoter across >800 tissues and cell types ^26^. For each gene with an average model performance of at least 0.05 as in ^4^ (*i.e.* a predictable gene), we counted the number of selected (and unselected) TF binding events in distal regulatory regions overlapping FANTOM5 enhancers associated to that gene. Individual gene statistics were aggregated, and the overall enrichment of selected features in enhancer regions was calculated using Fisher’s exact test.

### Validating the correlation between TF binding and gene expression in vivo

Next, for the key TF binding events identified by TF2Exp models, we sought to validate whether the TF binding correlates with the gene expression *in vivo*. We obtained CTCF ChIP-seq mapped data (BAM files) for 45 LCLs ^43^ and PU.1 for another set of 45 LCLs (38/45 overlap with CTCF LCLs) ^44^ from the 358 LCLs in the original training data. For each TF binding event, the TF binding signal was quantified as the number of ChIP-seq reads in each ChIP-seq experiment using HOMER ^45^. TF binding signals were then normalized through multiple steps, including scaling by library size, averaging between replicates of each individual, converting to standard deviation units (standardization), performing quantile normalization and removing batch effects by PEER ^37^. The resulting normalized data constitutes the *in vivo* TF binding signal for each TF binding event in each LCL.

We reserved the LCLs for which the extra ChIP-seq data was available as testing sets (one set for each of the two TFs). TF2Exp models were retrained on the non-testing LCLs, identifying 370 CTCF and 309 PU.1 TF binding events as key features for the subset of predictable genes. As less than 10% of *in vivo* TF binding events have been previously reported to show variable binding (greater inter-individual variance than intra-replicate variance)^46, 47^, we anticipated that the majority of selected TF binding events in the testing cases would be invariable. To focus on potential variable TF binding events (and minimize multiple testing impacts), we restricted the analysis to the subset of TF binding events which exhibit a strong DeepSEA score variance (top 10% of all TF binding events), resulting in 83 CTCF and 72 PU.1 selected TF binding events. Then, we assessed the correlation between the *in vivo* TF binding of the selected events and the associated gene expression in the two testing sets. Recognizing the small sample size, we estimated the minimum detectable correlation coefficient for the given testing size (n=45) at significance of 0.05 and power of 0.6 using the pwr package ^48^.

### External validation with microarray expression data

For external validation of TF2Exp models, we relied on microarray data reporting expression levels of 15,997 Ensembl genes for LCLs of 80 CEU, 87 Chinese (CHB), and 89 Japanese (JPT) individuals ^27^. For these individuals, variant data was retrieved from the 1000 Genomes Project. We applied the TF2Exp model to predict gene expression levels from potentially altered TF binding events based on the variant data, and compared these predictions with the gene expression levels reported from the microarray.

### Code and data availability

The code and model training results can be found at www.github.com/wqshi/TF2Exp. Multiple packages have been used for data processing and model training, including BEDTools ^49^, vcftools ^50^, caret ^51^ and ggplot2 ^52^.

## Acknowledgements

The authors thank the Wasserman lab members for helpful discussions, Miroslav Hatas for systems support, and Dora Pak for management support. In particular, the authors thank Rachelle Farkas and David Arenillas for thorough proofreadeing of this manuscript. This work was supported by Genome Canada/CIHR/Genome BC [174DE], the Natural Sciences and Engineering Research Council of Canada [RGPIN355532-10], the National Institutes of Health (USA) [1R01GM084875], and PhD fellowship from China Scholarship Council [201206110038] [to WS]. Funding has been provided by the BC Children’s Hospital Foundation to WWW. Funding for open access charge: Genome Canada Bioinformatics and Computational Biology (255ONT), Canadian Institutes of Health Research (CIHR): Bioinformatics and Computational Biology (BOP-149430).

## Authors’ contributions

With WWW, WS conceived the research and designed the study. WS conducted all the analyses with WWW and OF, and generated all figures and all tables. WS wrote the manuscript, which OF and WWW reviewed and revised.

## References

1. Hindorff LA, Sethupathy P, Junkins HA, Ramos EM, Mehta JP, Collins FS, et al. Potential etiologic and functional implications of genome-wide association loci for human diseases and traits. Proceedings of the National Academy of Sciences of the United States of America 2009, 106(23): 9362–9367.

2. Consortium EP. An integrated encyclopedia of DNA elements in the human genome. Nature 2012, 489(7414): 57–74.

3. Lappalainen T, Sammeth M, Friedlander MR, t Hoen PA, Monlong J, Rivas MA, et al. Transcriptome and genome sequencing uncovers functional variation in humans. Nature 2013, 501(7468): 506–511.

4. Manor O, Segal E. Robust prediction of expression differences among human individuals using only genotype information. PLoS genetics 2013, 9(3): e1003396.

5. Gamazon ER, Wheeler HE, Shah KP, Mozaffari SV, Aquino-Michaels K, Carroll RJ, et al. A gene-based association method for mapping traits using reference transcriptome data. Nature genetics 2015, 47(9): 1091–1098.

6. Farh KK, Marson A, Zhu J, Kleinewietfeld M, Housley WJ, Beik S, et al. Genetic and epigenetic fine mapping of causal autoimmune disease variants. Nature 2014.

7. Gibson G. Rare and common variants: twenty arguments. Nature reviews Genetics 2012, 13(2): 135–145.

8. Lappalainen T. Functional genomics bridges the gap between quantitative genetics and molecular biology. Genome research 2015, 25(10): 1427–1431.

9. Wasserman WW, Sandelin A. Applied bioinformatics for the identification of regulatory elements. Nature reviews Genetics 2004, 5(4): 276–287.

10. Zhou J, Troyanskaya OG. Predicting effects of noncoding variants with deep learning-based sequence model. Nature methods 2015, 12(10): 931–934.

11. Quang D, Xie X. DanQ: a hybrid convolutional and recurrent deep neural network for quantifying the function of DNA sequences. Nucleic acids research 2016.

12. Shi W, Fornes O, Mathelier A, Wasserman WW. Evaluating the impact of single nucleotide variants on transcription factor binding. Nucleic acids research 2016: gkw691.

13. Chen J, Rozowsky J, Galeev TR, Harmanci A, Kitchen R, Bedford J, et al. A uniform survey of allele-specific binding and expression over 1000-Genomes-Project individuals. Nature communications 2016, 7: 11101.

14. Cusanovich DA, Pavlovic B, Pritchard JK, Gilad Y. The functional consequences of variation in transcription factor binding. PLoS genetics 2014, 10(3): e1004226.

15. Spivakov M. Spurious transcription factor binding: non-functional or genetically redundant? BioEssays: news and reviews in molecular, cellular and developmental biology 2014, 36(8): 798–806.

16. Genomes Project C, Abecasis GR, Auton A, Brooks LD, DePristo MA, Durbin RM, et al. An integrated map of genetic variation from 1,092 human genomes. Nature 2012, 491(7422): 56–65.

17. Friedman J, Hastie T, Tibshirani R. Regularization Paths for Generalized Linear Models via Coordinate Descent. Journal of statistical software 2010, 33(1): 1–22.

18. McLean CY, Bristor D, Hiller M, Clarke SL, Schaar BT, Lowe CB, et al. GREAT improves functional interpretation of *cis*-regulatory regions. Nature biotechnology 2010, 28(5): 495–501.

19. Natarajan A, Yardimci GG, Sheffield NC, Crawford GE, Ohler U. Predicting cell-type-specific gene expression from regions of open chromatin. Genome research 2012, 22(9): 1711–1722.

20. Ong CT, Corces VG. CTCF: an architectural protein bridging genome topology and function. Nature reviews Genetics 2014, 15(4): 234–246.

21. Holwerda SJ, de Laat W. CTCF: the protein, the binding partners, the binding sites and their chromatin loops. Philosophical transactions of the Royal Society of London Series B, Biological sciences 2013, 368(1620): 20120369.

22. Lotem J, Levanon D, Negreanu V, Bauer O, Hantisteanu S, Dicken J, et al. Runx3 at the interface of immunity, inflammation and cancer. Biochimica et biophysica acta 2015, 1855(2): 131–143.

23. Hagman J, Ramirez J, Lukin K. B lymphocyte lineage specification, commitment and epigenetic control of transcription by early B cell factor 1. Current topics in microbiology and immunology 2012, 356: 17–38.

24. Iwafuchi-Doi M, Zaret KS. Pioneer transcription factors in cell reprogramming. Genes & development 2014, 28(24): 2679–2692.

25. Battle A, Mostafavi S, Zhu X, Potash JB, Weissman MM, McCormick C, et al. Characterizing the genetic basis of transcriptome diversity through RNA-sequencing of 922 individuals. Genome research 2014, 24(1): 14–24.

26. Andersson R, Gebhard C, Miguel-Escalada I, Hoof I, Bornholdt J, Boyd M, et al. An atlas of active enhancers across human cell types and tissues. Nature 2014, 507(7493): 455–461.

27. Stranger BE, Montgomery SB, Dimas AS, Parts L, Stegle O, Ingle CE, et al. Patterns of cis regulatory variation in diverse human populations. PLoS genetics 2012, 8(4): e1002639.

28. Taneera J, Fadista J, Ahlqvist E, Atac D, Ottosson-Laakso E, Wollheim CB, et al. Identification of novel genes for glucose metabolism based upon expression pattern in human islets and effect on insulin secretion and glycemia. Human molecular genetics 2015, 24(7): 1945–1955.

29. Pedersen HK, Gudmundsdottir V, Brunak S. Pancreatic Islet Protein Complexes and Their Dysregulation in Type 2 Diabetes. Frontiers in genetics 2017, 8: 43.

30. Consortium F, the RP, Clst, Forrest AR, Kawaji H, Rehli M, et al. A promoter-level mammalian expression atlas. Nature 2014, 507(7493): 462–470.

31. Wagner JR, Busche S, Ge B, Kwan T, Pastinen T, Blanchette M. The relationship between DNA methylation, genetic and expression inter-individual variation in untransformed human fibroblasts. Genome biology 2014, 15(2): R37.

32. Huang YF, Gulko B, Siepel A. Fast, scalable prediction of deleterious noncoding variants from functional and population genomic data. Nature genetics 2017, 49(4): 618–624.

33. Tjong H, Li W, Kalhor R, Dai C, Hao S, Gong K, et al. Population-based 3D genome structure analysis reveals driving forces in spatial genome organization. Proceedings of the National Academy of Sciences of the United States of America 2016, 113(12): E1663–1672.

34. Genomes Project C, Auton A, Brooks LD, Durbin RM, Garrison EP, Kang HM, et al. A global reference for human genetic variation. Nature 2015, 526(7571): 68–74.

35. Grubert F, Zaugg JB, Kasowski M, Ursu O, Spacek DV, Martin AR, et al. Genetic Control of Chromatin States in Humans Involves Local and Distal Chromosomal Interactions. Cell 2015, 162(5): 1051–1065.

36. Patro R, Mount SM, Kingsford C. Sailfish enables alignment-free isoform quantification from RNA-seq reads using lightweight algorithms. Nature biotechnology 2014, 32(5): 462–464.

37. Stegle O, Parts L, Piipari M, Winn J, Durbin R. Using probabilistic estimation of expression residuals (PEER) to obtain increased power and interpretability of gene expression analyses. Nature protocols 2012, 7(3): 500–507.

38. Aken BL, Ayling S, Barrell D, Clarke L, Curwen V, Fairley S, et al. The Ensembl gene annotation system. Database: the journal of biological databases and curation 2016, 2016.

39. Neph S, Vierstra J, Stergachis AB, Reynolds AP, Haugen E, Vernot B, et al. An expansive human regulatory lexicon encoded in transcription factor footprints. Nature 2012, 489(7414): 83–90.

40. Chang CC, Chow CC, Tellier LC, Vattikuti S, Purcell SM, Lee JJ. Second-generation PLINK: rising to the challenge of larger and richer datasets. GigaScience 2015, 4: 7.

41. Team RC. R: A language and environment for statistical computing. 2013.

42. Chatr-Aryamontri A, Breitkreutz BJ, Oughtred R, Boucher L, Heinicke S, Chen D, et al. The BioGRID interaction database: 2015 update. Nucleic acids research 2015, 43(Database issue): D470–478.

43. Ding Z, Ni Y, Timmer SW, Lee BK, Battenhouse A, Louzada S, et al. Quantitative Genetics of CTCF Binding Reveal Local Sequence Effects and Different Modes of X-Chromosome Association. PLoS genetics 2014, 10(11): e1004798.

44. Waszak SM, Delaneau O, Gschwind AR, Kilpinen H, Raghav SK, Witwicki RM, et al. Population Variation and Genetic Control of Modular Chromatin Architecture in Humans. Cell 2015, 162(5): 1039–1050.

45. Heinz S, Benner C, Spann N, Bertolino E, Lin YC, Laslo P, et al. Simple combinations of lineage-determining transcription factors prime cis-regulatory elements required for macrophage and B cell identities. Molecular cell 2010, 38(4): 576–589.

46. Kasowski M, Grubert F, Heffelfinger C, Hariharan M, Asabere A, Waszak SM, et al. Variation in transcription factor binding among humans. Science 2010, 328(5975): 232–235.

47. Kasowski M, Kyriazopoulou-Panagiotopoulou S, Grubert F, Zaugg JB, Kundaje A, Liu Y, et al. Extensive variation in chromatin states across humans. Science 2013, 342(6159): 750–752.

48. Champely S. pwr: Basic functions for power analysis. R package version 2012, 1(1).

49. Quinlan AR. BEDTools: The Swiss-Army Tool for Genome Feature Analysis. Current protocols in bioinformatics / editoral board, Andreas D Baxevanis [et al] 2014, 47: 11 12 11–34.

50. Danecek P, Auton A, Abecasis G, Albers CA, Banks E, DePristo MA, et al. The variant call format and VCFtools. Bioinformatics 2011, 27(15): 2156–2158.

51. Kuhn M. caret: Classification and Regression Training. 2015.

52. Wickham H. ggplot2: elegant graphics for data analysis. Springer Science & Business Media, 2009.

